# The metalloproteinase inhibitor Prinomastat reduces AML growth, prevents stem cell loss and improves chemotherapy effectiveness

**DOI:** 10.1101/2020.12.01.393157

**Authors:** Chiara Pirillo, Myriam Haltalli, Sara Gonzalez Anton, Valentina Tini, Isabella Kong, Edwin Hawkins, Brunangelo Falini, Andrea Marra, Delfim Duarte, Cristina Lo Celso

## Abstract

Acute myeloid leukemia (AML) is a blood cancer of the myeloid lineage. Its prognosis remains poor, highlighting the need for new therapeutic and precision medicine approaches. AML symptoms often include cytopenias, linked to loss of healthy hematopoietic stem and progenitor cells (HSPCs). The mechanism behind HSPC decline is complex and still poorly understood. Here, intravital microscopy (IVM) of a well-established experimental model of AML allows direct observation of the interactions between healthy and malignant cells in the bone marrow (BM), suggesting that physical dislodgment of healthy cells by AML through damaged vasculature may play an important role. Numerous human leukemia types, particularly MLL-AF9 samples, show high expression levels of multiple matrix metalloproteinases (MMPs). Therefore, we evaluate the therapeutic potential of the MMP inhibitor (MMPI) prinomastat. IVM analyses of treated mice reveal reduced vascular permeability and healthy cell clusters in circulation, and lower AML cell speed. Furthermore, treated mice have decreased BM infiltration, increased retention of healthy HSPCs in the BM and increased survival following chemotherapy. Overall, our results suggest that MMPIs could be a promising complementary therapy to reduce AML growth and limit the loss of HSPC and BM vascular damage caused by MLL-AF9 and possibly other AML subtypes.

## Introduction

Acute myeloid leukemia (AML) is the most common form of acute leukemia in adults. While progress has been made in treatment development, relapse incidence remains high, resulting in poor prognosis^1^. One challenge with developing effective AML treatments is the extensive diversity in the disease biology, underpinned by the large number of genetic alterations driving disease development^2^. Precision medicine is therefore a sought-after approach to tackle AML successfully. AML patients develop severe cytopenia due to loss of healthy hematopoietic stem and progenitor cells (HSPCs) driven by remodeling of the bone marrow (BM) microenvironment by malignant cells. Healthy blood cell count recovery following chemotherapy has emerged as the second most important predictor of disease-free survival after minimal residual disease^3,4^. It critically depends on healthy HSPCs driving hematopoietic recovery from the BM, therefore therapeutic interventions that can strengthen healthy hematopoiesis and protect HSPCs’ microenvironments are necessary.

Well-established murine experimental models of AML allow studying the competition between healthy and AML cells within the BM, and have shown a clear anti-correlation between the number of healthy HSPCs and AML cells^5–7^. Understanding the principles of healthy vs. malignant cells’ competition is important to identify novel therapeutic targets. We and others have reported absence of widespread apoptosis of healthy cells^5,7^, increased size and stiffness of leukemia cells^8^ and increased vascular permeability in the BM of AML-burdened mice^9^, raising the hypothesis that AML cells may orchestrate a physical displacement of healthy cells. Extracellular matrix (ECM) surrounds and supports all cells, including hematopoietic precursors in the BM^10^.

Dysfunction in ECM remodeling was demonstrated to facilitate solid cancer invasion and metastasis^11^. Its role in leukemia is still understudied, however it has been reported that matrix stiffness can affect leukemia growth and that myeloid malignant cell proliferation is enhanced on softer matrices^12^. Intravital microscopy (IVM) uniquely enables direct observation of the interactions between AML and healthy hematopoietic cells within the BM microenvironment^13^. With it, here we identify increased vascular permeability earlier than previously reported, and abnormal egress of healthy cells into circulation. Deregulated expression of matrix metalloproteinases (MMPs) in murine and human AML samples suggests that these enzymes may contribute to the processes observed. Using the metalloproteinase inhibitor (MMPI) prinomastat (AG3340), we identify MMPs as a promising target to inhibit AML progression, protect BM vasculature, retain BM HSPCs and improve chemotherapy efficacy.

## Combined Results and Discussion

To directly observe competition between AML and healthy hematopoietic cells, we generated chimeric mice bearing non-fluorescent stroma and membrane-bound mTomato^+^ hematopoietic cells, injected them with YFP^+^ MLL-AF9 AML blasts and examined the calvarium BM by IVM (Figure 1). When we compared AML-burdened and healthy chimeras, we detected exclusively in leukemic mice multiple non-malignant mTomato^+^ cells intravasating as clusters (Figure 1A and Supplementary Video 1). Consistent with this, while we observed mostly single cells and occasionally doublets in circulation in healthy mice, we detected clusters of up to eight healthy cells circulating in mice with intermediate leukemia infiltration (10-25% blasts in peripheral blood (PB); Figure 1B-C and Supplementary Video 2). Of note, these circulating clusters were no longer frequent once BM was fully infiltrated and healthy hematopoiesis outcompeted. Moreover, longitudinal flow cytometry analysis of the PB of AML-burdened mice highlighted increasing numbers of undifferentiated (Lineage^−^) healthy cells in circulation. These were directly proportional to BM AML infiltration and followed the inverse trajectory of Lineage^−^ cells’ abundance in the BM (Supplementary Figure 1). These data indicate that healthy cells are displaced from the BM parenchyma into circulation during AML growth and provide a mechanism for healthy cells’ ousting from BM. Given these observations, and because AML cells are known to grow in foci where BM stroma is locally remodeled^13^, we hypothesized that healthy cell displacement may result from a combination of microenvironment disruption and physical displacement towards vessels. We selected matrix metalloproteinases (MMPs) as a likely candidate driver of this mechanism because they are involved in ECM remodeling^11,14^, ECM alterations are important in solid cancer progression and invasion^11,15^, and MMPs have altered expression and/or function in several cancers^16^, including leukemia^17–19^. Moreover, MMPs contribute to regulating vascular permeability^20^ and increased BM vascular leakiness in mice fully infiltrated by AML has been reported^9^. Interestingly, we could measure increased BM vascular leakiness in mice already at early disease stages (<10% PB infiltration) (Figure 1D and E and Supplementary Video 3). This suggested that early vascular leakiness may contribute to the observed intravasation of healthy cell clusters. When we analyzed a transcriptomic dataset that we previously generated^13^, we identified that murine AML cells express significantly higher levels of multiple MMPs compared to healthy granulocyte/monocyte progenitors (GMPs), which they derive from and resemble to phenotypically (Figure 1F-G). Importantly, analysis of the Cancer Genome Atlas (TCGA) database identified deregulation of MMPs’ expression across AML subtypes, with the MLL-AF9 AML variant showing upregulation of most MMPs, particularly MMP2, 9 and 14 (Figure 1H).

**Figure 1.**
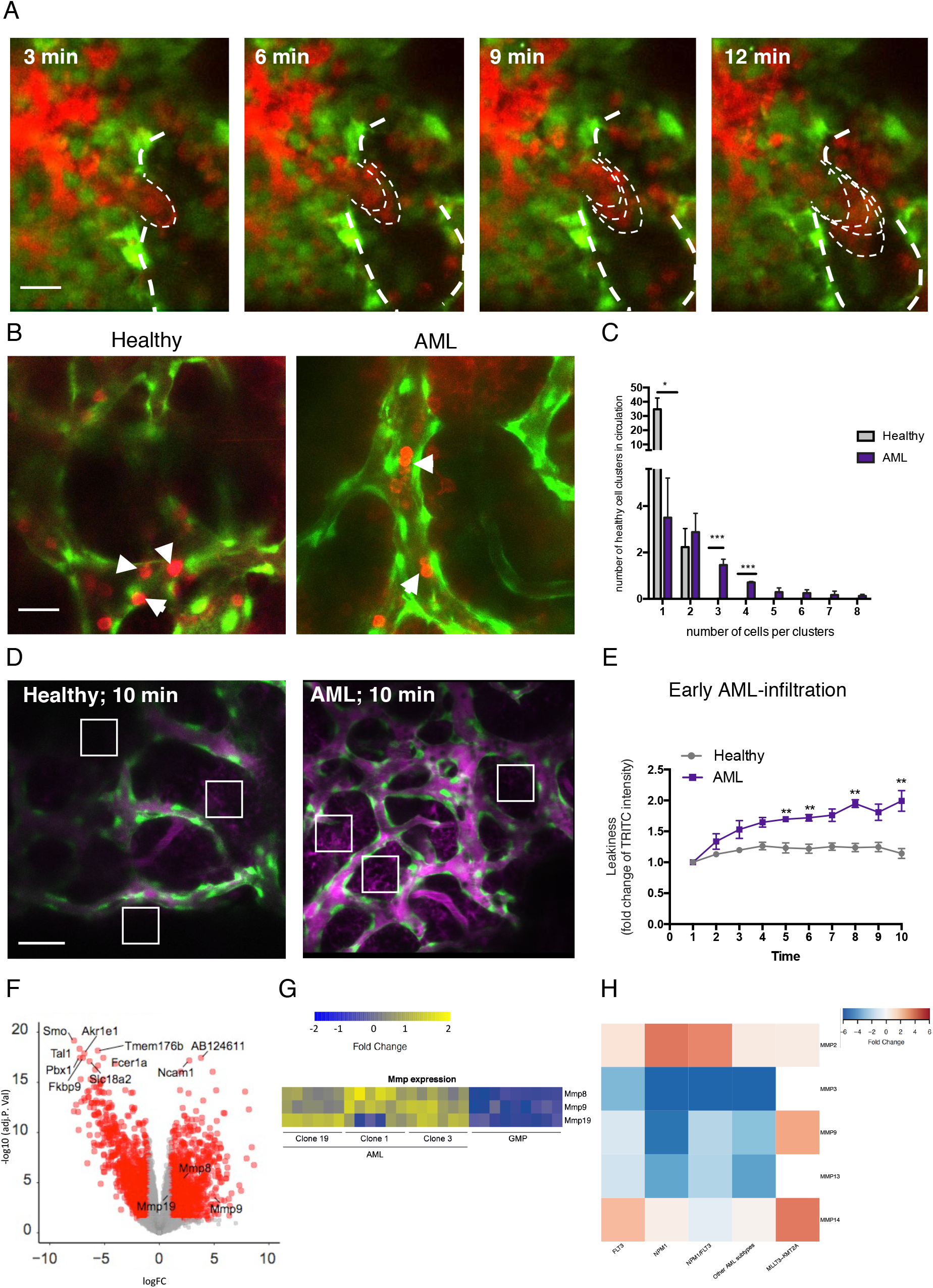
Healthy haematopoietic cells are dislodged from the BM. (A) Selected frames from representative time-lapse IVM of calvarium BM in AML-burdened Flk1-GFP mice, showing endothelial cells (GFP^+^, brightest green, thick dotted lines highlight vessel inner edge), YFP^+^ AML cells (dimmer green), and mTomato^+^ healthy haematopoietic cells (red) egressing from the BM as a cluster (thin dotted lines show cluster leading edge protruding into the vessel over time). (B) Selected frames from representative time-lapse IVM of calvarium BM in healthy (left) and AML-burdened (right) Flk1-GFP mice. Green: ECs. Red: mTomato^+^ healthy hematopoietic cells. Arrows point at circulating healthy cell clusters. (C) Quantification of circulating healthy single cells and clusters in healthy and AML-burdened mice. Data representative of and pooled from 8 mice from two independent experiments. (D) Vascular leakiness assessed by time-lapse IVM of randomly selected regions (white boxes) within the calvarium of Flk1-GFP mice following administration of 3mg TRITC-dextran. Selected representative frames of healthy and leukemic (<10% AML blood infiltration) mice are shown. (E) Quantification of the fold change in TRITC-dextran intensity in the BM parenchyma of healthy and early infiltrated leukemic mice. *n* = 3 control and 3 leukemic mice, 3 areas measured/mouse. (F) Volcano plot for DEGs between AML and GMPs with cut off value logFC≥1 or log FC≤1 and pvalue≤0.05. Top 10 DEGs and MMPs are labelled (G) Gene expression heatmap for MMP-8, MMP-9 and MMP-19 for AML and GMP. *n* = 9 control and *n* = 9 leukemic mice. (H) Heatmap of Prinomastat-targeted MMPs from TCGA RNA-seq data from healthy and AML adult patients. MMP-9 and MMP-14 are specifically upregulated within *MLLt3-KMT2A* AML that is most similar to the MLL-AF9 murine AML model. Red and blue: up- and downregulation, respectively. All scale bars represent 80μm. All data are mean ± s.e.m. * p < 0.05; ** p < 0.01; *** p < 0.001. *P* values determined by multiple *t*-tests with post-hoc Holm-Sidak corrections

Next, we tested the effect of treating mice with the selective MMP inhibitor (MMPI) prinomastat (AG3340), which targets multiple MMPs including MMP9, which we had found upregulated in both murine and human AML cells. Mice were administered prinomastat or PBS daily for two weeks starting from day 7 post-AML blasts injection, and were analyzed during and at the end of treatment (Figure 2A). Prinomastat-treated mice had fewer and smaller healthy cell clusters and more single cells in circulation than PBS-treated mice (Figure 2B - C and Supplementary Videos 4 - 5). Moreover, vascular leakiness was significantly reduced (Figure 2D - E and Supplementary Video 6), with prinomastat-treated mice showing dextran extravasation rates similar to those of healthy controls (see Figure 1E). This suggested that prinomastat treatment rescues vascular leakiness in the context of AML, and this correlates with reduced intravasation of cell clusters. Prinomastat treatment reduces VEGF levels in solid cancers^21^. AML cells produce VEGF, leading to failed angiogenesis^13^ and increased vascular permeability^9^. In our experiments prinomastat did not affect VEGF levels in BM tissue (Figure 2F), nor endothelial cell (EC) numbers; however, it reduced Reactive Oxygen Species (ROS) in ECs, a factor recently linked to increased BM vascular leakiness^9,22^ (Figure 2G). It has been suggested that ECs express MMPs and remodel local ECM in response to angiogenic stimuli^20^, and the question remains open whether the reduced vascular permeability observed here is due to inhibition of AML-derived, EC-derived MMPs, or both.

**Figure 2.**
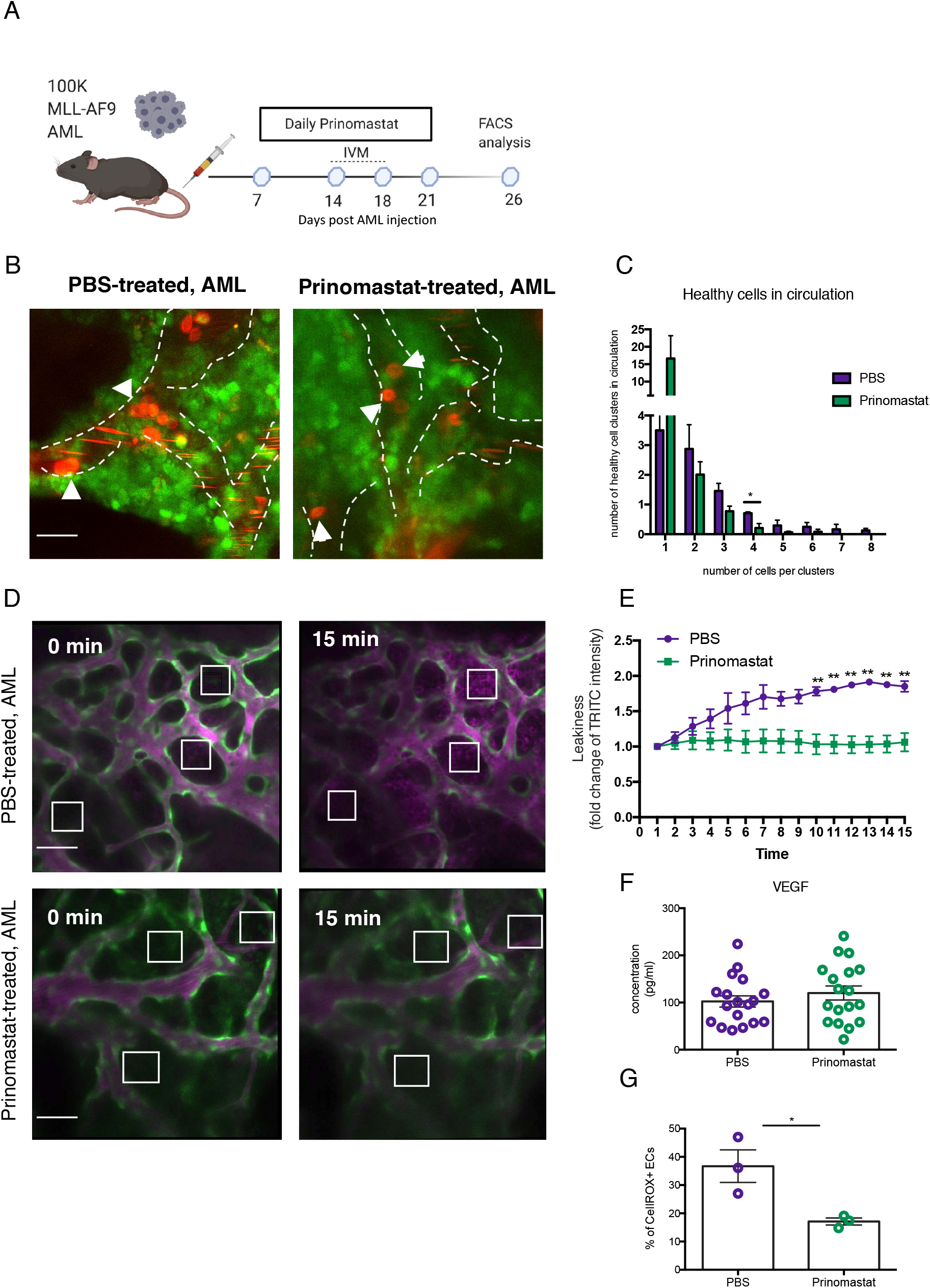
Prinomastat reduces circulating healthy cell clusters and rescues vascular leakiness. (A) Schematic of the prinomastat treatment regime adopted. (B) Selected, representative time-frames of time-lapse IVM showing YFP^+^ AML blasts (green) and circulating mTomato^+^ healthy haematopoietic cells (red) in PBS- and prinomastat-treated leukemic mice. Dashed white lines delineate vessel walls. Arrowheads point at clusters of healthy cells in PBS-treated and single cells in prinomastat-treated leukemic mice, respectively. Scale bar represents 25μm. (C) Quantification of circulating healthy single cells and clusters in PBS- and prinomastat-treated leukemic mice. * p < 0.05. Data pooled from 6 mice in total from two independent experiments. (D) Vascular leakiness assessed by time-lapse IVM of randomly selected regions (white boxes) within the calvarium of Flk1-GFP following administration of TRITC-dextran i.v. Selected representative frames from PBS- and prinomastat-treated leukemic mice are shown. Scale bar represents 80μm. (E) Quantification of the fold change in TRITC-dextran intensity in the BM parenchyma of PBS- and prinomastat-treated mice. *n* = 3 PBS- and *n* = 3 prinomastat-treated, leukemic mice. (F) VEGF levels (pg/mL) measured in the BM supernatant of PBS- and prinomastat-treated leukemic mice by ELISA. *n* = 18 mice per condition (G) Quantification of CellROX^+^ endothelial cells (CD45^−^/Ter119^−^ CD31^+^ Sca-1^+^) in PBS- and prinomastat-treated mice at late time point. n= 3 per condition. All data are mean ± s.e.m.* p < 0.05; *** p < 0.001; **** p < 0.0001. *P* values determined by multiple *t*-tests with post-hoc Holm-Sidak corrections

Next, we asked whether prinomastat affects AML cells. IVM tilescans indicated the BM of prinomastat-treated mice contained fewer leukemic cells (Figure 3A), and reduced BM infiltration was confirmed by flow cytometry analysis of long bones (Figure 3B). This was linked to reduced AML cell proliferation and increased apoptosis (Figure 3C and D). Unexpectedly, AML blast infiltration in the PB of prinomastat- and PBS-treated animals was similar (Figure 3E). This could be explained by a reduced ability of circulating AML cells to extravasate and re-enter the BM parenchyma and would be consistent with the observed reduced vascular permeability and with reduced spleen infiltration (Supplementary Figure 2), which in this model is driven by trafficking of cells from BM^7^. Finally, time-lapse IVM imaging revealed that the average speed of AML cells within the BM parenchyma was significantly reduced in prinomastat-treated mice (Figure 3F). In particular, while most AML cells exhibited a migratory behavior^23^, in untreated animals we would consistently observe a significant proportion of AML cells with a complex and highly dynamic morphology, which we refer to as ‘explorative cells’ because we often found them moving between healthy cells (Figure 3G and Supplementary Video 7). These explorative cells were typically faster and exhibited a more directional movement than the remaining bulk of AML cells, which were round and slower (Figure 3H - K). Interestingly, the number of explorative cells was significantly reduced in prinomastat-treated mice (Figure 3L). Altogether, our data demonstrate that prinomastat treatment affects not only BM vascular leakiness but also the leukemia cells.

**Figure 3.**
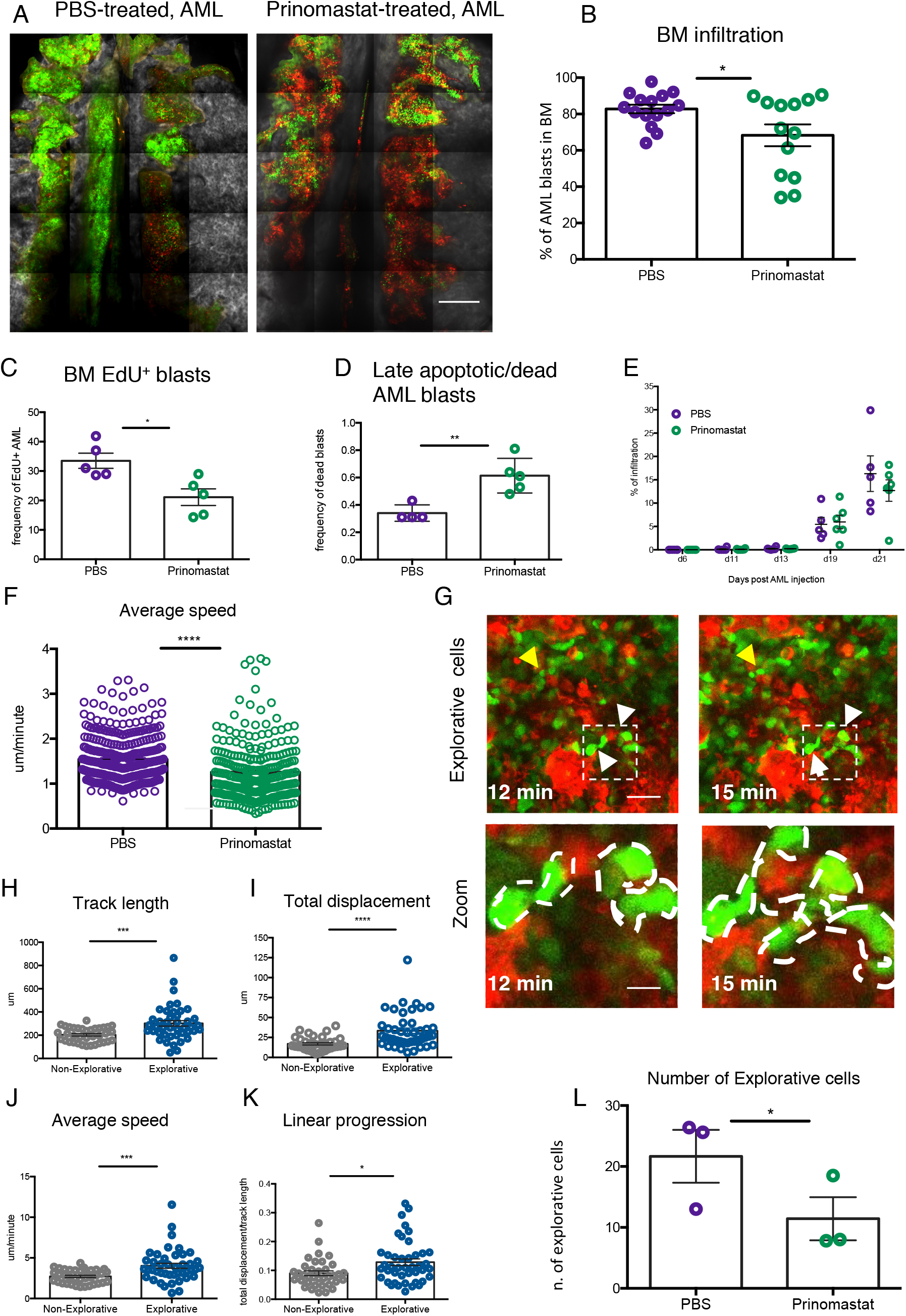
Multiple effects of prinomastat on leukemic blasts in the BM. (A) Representative tilescan maximum projections of calvarium BM of PBS- and prinomastat-treated AML-burden mice. Red: mTomato^+^ healthy haematopoietic cells; green: YFP^+^ AML blasts; grey: bone collagen. Scale bar represents 500 μm. (B) AML blast infiltration in the long bones of PBS- and prinomastat-treated leukemic mice. Each dot represents one mouse. *n* = 15 PBS-treated and *n* = 13 Prinomastat-treated mice pooled from 2 independent experiments. (C) Proportion of proliferating (EdU^+^) and (D) apoptotic/dead (Annexin V+) AML blasts in the BM of PBS- and prinomastat-treated mice. *n =* 4 and 5 mice per group, respectively. (E) Percentage of AML blasts measured in the peripheral blood of PBS- and Prinomastat-treated mice throughout disease progression. *n* = 6 mice per group (F) Average speed of AML blasts in calvarium BM of PBS- and prinomastat-treated AML-burdened mice. *n* = 338 and *n* = 293 cells per group, pooled from n=3 mice per group from 2 independent experiments. (G) Selected frames from representative time-lapse IVM of BM calvarium with intermediate infiltration. Red: mTomato^+^ healthy haematopoietic cells; green: YFP^+^ AML blasts. White and yellow arrows: examples of explorative and non-explorative cells, respectively. Boxed areas are at higher magnification below. Scale bars represent 80μm and 25μm respectively. (H) Quantification of track length, (I) total displacement, (J) average speed and (K) linear progression of non-explorative and explorative cells *n =* 68 explorative cells and *n* = 60 non-explorative cells pooled from 3 mice in two independent experiments. (L) Number of explorative cells in PBS- and prinomastat-treated, leukemic mice in the calvarium BM. *n* = 3 mice per group. All data are mean ± s.e.m. *p < 0.05; *** p < 0.001; **** p < 0.0001. *P* values determined by student *t*-tests

Given the effects of prinomastat on both vasculature and AML cells, we hypothesized that the MMPI may have a protective effect on residual HSPC populations too. We therefore grouped prinomastat- and PBS-treated mice in two categories, mid- and late-infiltrated, based on measured BM AML burden (40-75% and 75-100% blasts in BM, respectively), and assessed the number of residual BM HSPCs. In each category, prinomastat-treated mice showed significantly higher numbers of residual long-term HSCs (phenotypically defined as Lineage^−^c-Kit^+^Sca-1^+^[LKS]CD48^−^CD150^+^), and multipotent progenitor cells (LKS CD48^+^CD150^−^), while short-term HSCs (LKS CD48^−^ CD150^−^) numbers were more variable (Figure 4A – C). HSPC maintenance within the marrow is mediated by multiple niche-derived molecules, with CXCL12 and SCF playing pivotal roles^24^; however, while both cytokines showed decreased levels in AML-burdened mice, their levels remained low in prinomastat-treated mice (Figure 4D and E). Together, our data suggest that the positive effect of prinomastat on the retention of HSPCs in their natural microenvironments is likely mediated by a combination of the effects on ECM remodeling/vasculature permeability and blasts’ dynamics, rather than on changes to the cytokine milieu.

**Figure 4:**
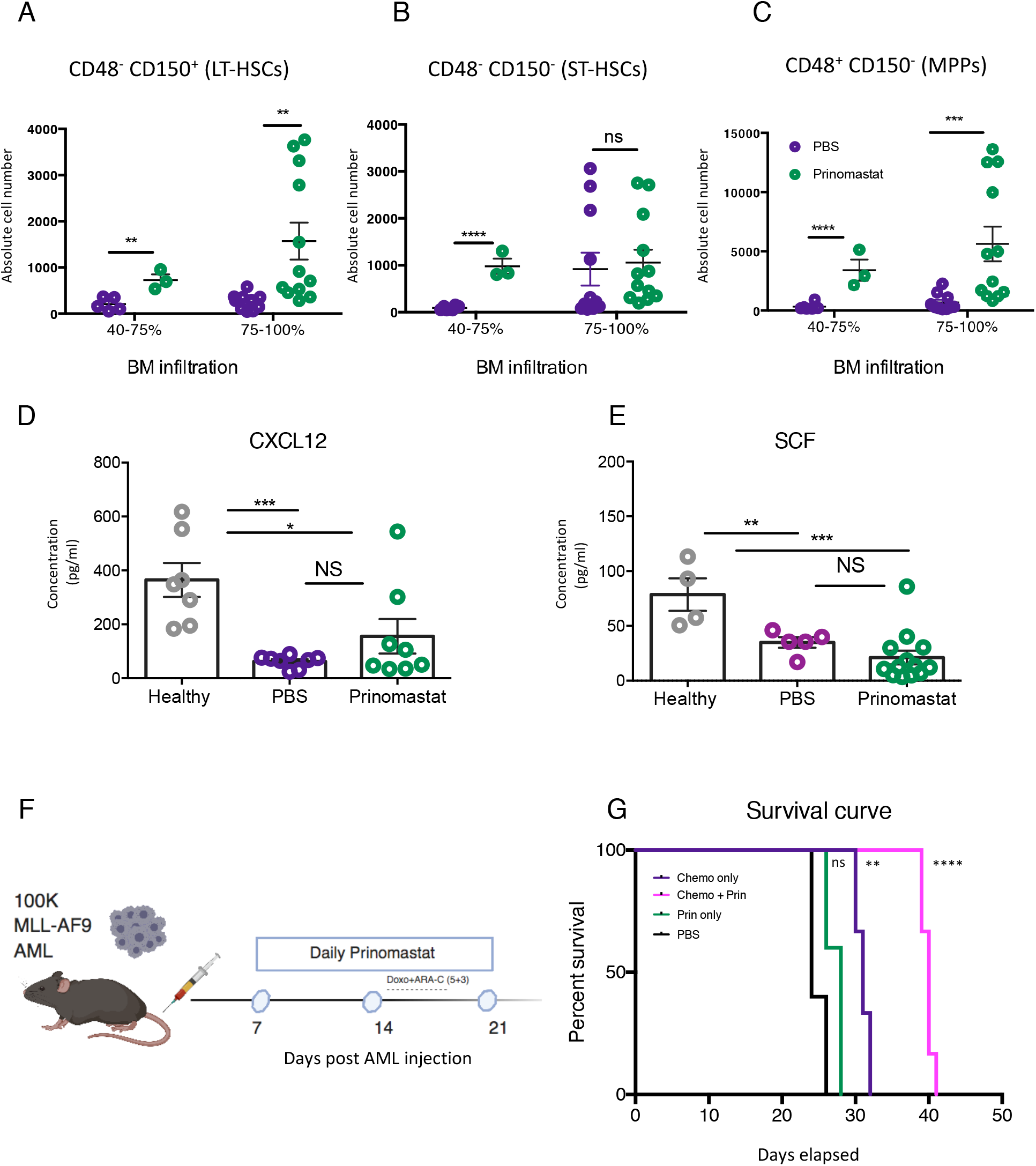
Prinomastat protects healthy HSPCs and, when combined with chemotherapy, increases survival. The absolute number of (A) (LKS CD48^−^CD150^+^), (B) (LKS CD48^−^ CD150^−^) and (C) (LKS CD48^+^CD150^−^) analysed in the BM of PBS- and prinomastat-treated, leukemic mice throughout disease progression. Each dot represents one mouse. *n* = at least 3 per infiltration point pooled from 3 independent experiments. (D) CXCL12 and (E) SCF levels (pg/mL) measured in BM supernatant from healthy control and PBS- and prinomastat-treated leukemic mice by ELISA. Each dot represents one mouse. *n* = 23 for CXCL12 and *n* = 22 for SCF, respectively. (F) Schematic of the combined prinomastat and chemotherapy treatment regime adopted. (G) Kaplan-Meyer curve showing survival of mice receiving chemotherapy alone, prinomastat alone or prinomastat in combination with chemotherapy or vehicle control (PBS). *n* = 6 for chemotherapy only and chemotherapy + prinomastat groups. *n* = 5 for prinomastat and PBS only groups. Data are shown as mean ± s.e.m. * p < 0.05; ** p < 0.01; *** p < 0.001; **** p < 0.0001; NS, not significant. *P* values determined using multiple *t*-tests with post-hoc Holm-Sidak corrections in (A – E) and the Log-rank test in (G)

Finally, we questioned whether prinomastat could improve AML outcome in a clinically relevant experimental setting. We therefore evaluated whether prinomastat administered alongside a conventional chemotherapy regime could improve animal survival. Mice were treated with prinomastat as described above, and chemotherapy was administered when PB infiltration reached 15 - 18% and following a well-established and clinically relevant regimen^13^ (Figure 4F). PBS, prinomastat-only and chemotherapy-only AML-burdened control groups were included in the experiment. Despite reducing leukemia infiltration, prinomastat treatment alone failed to prolong animal survival compared with the PBS- and chemotherapy-only control groups. However, when prinomastat and chemotherapy were administered in combination, we observed a significant increase in animal survival (Figure 4G). This finding suggests that prinomastat or other MMPIs could complement conventional AML treatment regimens to improve efficacy and overall survival, and is consistent with other reports indicating that ECs are promising targets for therapeutic intervention^9,13,25^ in leukaemia, that vascular ‘normalization’ is an important strategy when tackling multiple types of cancer^26^, and that leukemia cells grow less on stiffer, supposedly less degraded, matrices^12^.

The use of MMPIs in cancer therapy has been investigated in solid tumors with some controversial results^27^ and prinomastat reduced tumor burden and metastasis in pancreatic ductal adenocarcinoma and lung cancer, including in synergy with chemotherapy^28,29^.Overall, our work indicates that MMPs are deregulated in AML, and especially in the human MLL-AF9 subtype, which could especially benefit from MMPI treatment. Consistent with this, the MMPI prinomastat hindered disease progression and its effects on steady-state hematopoiesis in the well-established MLL-AF9 murine AML experimental model. AML cell proliferation, apoptosis and migratory ability were comprehensively affected resulting in lower BM infiltration. Simultaneously, prinomastat treatment rescued vascular leakiness and reduced ECs’ ROS, both linked to HSPCs intravasation^9,22^. Consistent with this, a higher number of residual HSPCs was retained in the BM. Furthermore, the combinatory use of prinomastat and conventional chemotherapy led to significantly increased survival in our experimental model. We propose that MMP inhibition should be further explored as a promising complementary therapeutic approach for AML patients with high MMPs expression levels and in combination with conventional induction/consolidation therapy and other more recently developed regimens.

## Methods

Additional methods and associated references are available in the Supplemental Methods.

### Study approval

All animal work was in accordance with the animal ethics committee (AWERB) at Imperial College London and UK Home Office regulations (ASPA, 1986).

### Statistics

GraphPad Prism was used for statistical analysis. Data are represented as mean ± SEM. Group means were compared using the unpaired Student’s t test. For multiple comparisons, multiple *t*-tests with post-hoc Holm-Sidak corrections or Bonferroni correction was used. A p value of less than 0.05 was considered significant.

## Author contributions

C.P. and C.L.C. conceived the project and contributed to the experimental designing. C.P. conducted the core experiments and data analysis. M.H. contributed to animal and flow cytometry experiments. S.G.A. contributed to flow cytometry experiments and analysis. A.M., V.T. and B.F. carried out the analysis on available data on Genome Atlas database for the MMPs expression profile for human data. D.D., I.K. and E.D.H. identified MMP expression in murine blasts. C.P. and C.L.C. led the writing of the manuscript with input from all authors.

## Acknowledgments

We thank staff of the core facilities at Imperial college London (Flow Cytometry, CBS facility, Healthcare NHS Trust) for their valuable help. This work was supported by Bloodwise (Gordon Piller PhD studentship to C.P.), Cancer Research UK (Programme Foundation award to C.L.C. and PhD studentship to S.G.A.), the Wellcome trust (PhD studentship PhD studentship 105398/Z/14/Z to M.H.), Associazione Italiana per la Ricerca sul Cancro (AIRC) to B.F. Treatment regime schematics were created with Biorender.com

## References

1. Longo DL, Döhner H, Weisdorf DJ, Bloomfield CD. Acute Myeloid Leukemia. New Engl J Medicine. 2015;373(12):1136–1152.

2. Network TCGAR. Genomic and Epigenomic Landscapes of Adult De Novo Acute Myeloid Leukemia. New Engl J Med. 2013;368(22):2059–2074. doi:10.1056/nejmoa1301689

3. Etienne A, et al. Impact of CRi on the Outcome of Elderly Patients with Untreated Acute Myeloid Leukemia (AML). Blood. 2008;112(11):2988–2988.

4. Chen X, et al. Relation of clinical response and minimal residual disease and their prognostic impact on outcome in acute myeloid leukemia. J Clin Oncol Official J Am Soc Clin Oncol. 2015;33(11):1258–1264.

5. Miraki-Moud F, et al. Acute myeloid leukemia does not deplete normal hematopoietic stem cells but induces cytopenias by impeding their differentiation. P Natl Acad Sci Usa. 2013;110(33):13576–13581.

6. Cheng H, et al. Leukemic marrow infiltration reveals a novel role for Egr3 as a potent inhibitor of normal hematopoietic stem cell proliferation. Blood. 2015;126(11):1302–1313.

7. Akinduro O, et al. Proliferation dynamics of acute myeloid leukaemia and haematopoietic progenitors competing for bone marrow space. Nat Commun. 2018;9(1):519.

8. Rosenbluth MJ, Lam WA, Fletcher DA. Force Microscopy of Nonadherent Cells: A Comparison of Leukemia Cell Deformability. Biophys J. 2006;90(8):2994–3003.

9. Passaro D, et al. Increased Vascular Permeability in the Bone Marrow Microenvironment Contributes to Disease Progression and Drug Response in Acute Myeloid Leukemia. Cancer Cell. 2017;32(3):324–341.e6.

10. Gattazzo F, Urciuolo A, Bonaldo P. Extracellular matrix: A dynamic microenvironment for stem cell niche. Biochimica Et Biophysica Acta Bba - Gen Subj. 2014;1840(8):2506–2519.

11. Lu P, Takai K, Weaver VM, Werb Z. Extracellular matrix degradation and remodeling in development and disease. Csh Perspect Biol. 2011;3(12):a005058–a005058.

12. Shin J-W, Mooney DJ. Extracellular matrix stiffness causes systematic variations in proliferation and chemosensitivity in myeloid leukemias. Proc National Acad Sci. 2016;113(43):12126–12131.

13. Duarte D, et al. Inhibition of Endosteal Vascular Niche Remodeling Rescues Hematopoietic Stem Cell Loss in AML. Cell Stem Cell. 2017;22(1):64–77.e6.

14. Daley WP, Peters SB, Larsen M. Extracellular matrix dynamics in development and regenerative medicine. J Cell Sci. 2008;121(3):255–264.

15. Lu P, Weaver VM, Werb Z. The extracellular matrix: a dynamic niche in cancer progression. J Cell Biology. 2012;196(4):395–406.

16. Gobin E, et al. A pan-cancer perspective of matrix metalloproteases (MMP) gene expression profile and their diagnostic/prognostic potential. Bmc Cancer. 2019;19(1):581.

17. Lin L-I, Lin D-T, Chang C-J, Lee C-Y, Tang J-L, Tien H-F. Marrow matrix metalloproteinases (MMPs) and tissue inhibitors of MMP in acute leukaemia: potential role of MMP-9 as a surrogate marker to monitor leukaemic status in patients with acute myelogenous leukaemia. Brit J Haematol. 2002;117(4):835–841.

18. Kamiguti AS, et al. The role of matrix metalloproteinase 9 in the pathogenesis of chronic lymphocytic leukaemia. Brit J Haematol. 2004;125(2):128–140.

19. Paupert J, Mas VM-D, Demur C, Salles B, Muller C. Cell-surface MMP-9 regulates the invasive capacity of leukemia blast cells with monocytic features. Cell Cycle. 2008;7(8):1047–1053.

20. Taraboletti G, D’Ascenzo S, Borsotti P, Giavazzi R, Pavan A, Dolo V. Shedding of the Matrix Metalloproteinases MMP-2, MMP-9, and MT1-MMP as Membrane Vesicle-Associated Components by Endothelial Cells. Am J Pathology. 2002;160(2):673–680.

21. Woenne EC, et al. MMP inhibition blocks fibroblast-dependent skin cancer invasion, reduces vascularization and alters VEGF-A and PDGF-BB expression. Anticancer Res. 2010;30(3):703–711.

22. Itkin T, et al. Distinct bone marrow blood vessels differentially regulate haematopoiesis. Nature. 2016;532(7599):323–328.

23. Duarte D, et al. Defining the in vivo characteristics of acute myeloid leukemia cells behavior by intravital imaging. Immunol Cell Biol. 2018;97(2):229–235.

24. Morrison SJ, Scadden DT. The bone marrow niche for haematopoietic stem cells. Nature. 2014;505(7483):327–334.

25. Barbier V, et al. Endothelial E-selectin inhibition improves acute myeloid leukaemia therapy by disrupting vascular niche-mediated chemoresistance. Nat Commun. 2020;11(1):2042.

26. Goel S, et al. Normalization of the Vasculature for Treatment of Cancer and Other Diseases. Physiol Rev. 2011;91(3):1071–1121.

27. Winer A, Adams S, Mignatti P. Matrix Metalloproteinase Inhibitors in Cancer Therapy: Turning Past Failures Into Future Successes. Mol Cancer Ther. 2018;17(6):1147–1155.

28. Alves F, et al. Inhibitory effect of a matrix metalloproteinase inhibitor on growth and spread of human pancreatic ductal adenocarcinoma evaluated in an orthotopic severe combined immunodeficient (SCID) mouse model. Cancer Lett. 2001;165(2):161–170.

29. Liu J, et al. Early combined treatment with carboplatin and the MMP inhibitor, prinomastat, prolongs survival and reduces systemic metastasis in an aggressive orthotopic lung cancer model. Lung Cancer. 2003;42(3):335–344.

