## Supplementary materials for "The metalloproteinase inhibitor Prinomastat reduces AML growth, prevents stem cell loss and improves chemotherapy effectiveness"



The authors have declared that no conflict of interest exists.

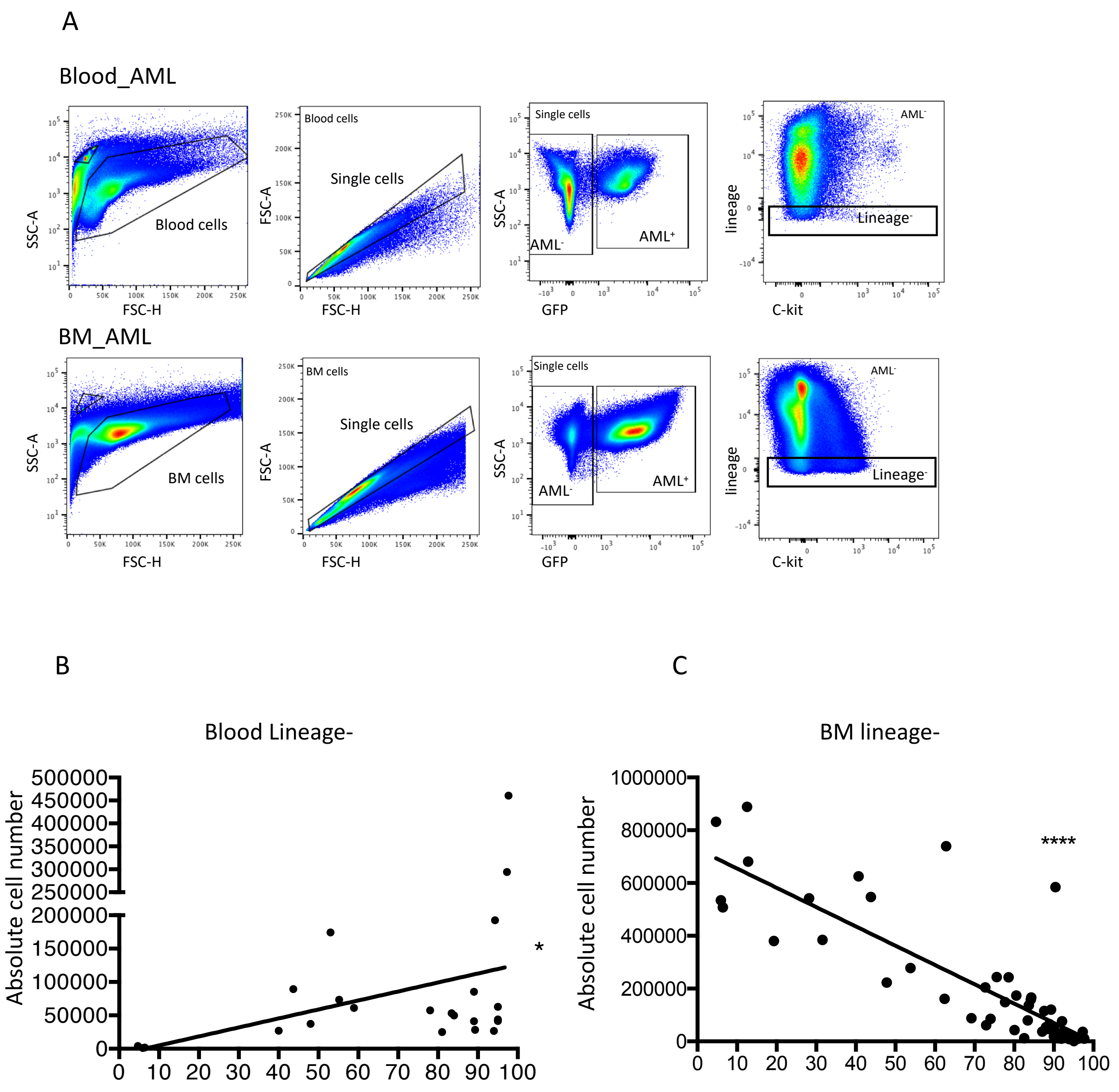


**Figure S1: Lineage negative cells increase in peripheral blood while decreasing in the BM in an AML-dependent manner.**

(A) Representative FACS plots from the blood and BM of leukemic mice analysed by flow cytometry. Black boxes indicate the gates used for gating on blood and BM cells to exclude debris, excluding doublets by gating on single cells, gating on $GFP^{+}$ AML to determine the infiltration and $GFP^{-}$ cells to then gating on the $Lineage^{-}$ population.

(B) Shows the increase in the absolute number of $Lineage^{-}$ cells in the blood that positively correlates with BM infiltration. n = 24 mice pooled from 2 independent experiments.

(C) The increase in BM infiltration negatively correlates to the absolute number of the $Lineage^{-}$ population within the BM. n = 57 mice pooled from 5 independent experiments.

*P* values determined using linear regression analysis, * $p < 0.05$; **** $p < 0.0001$

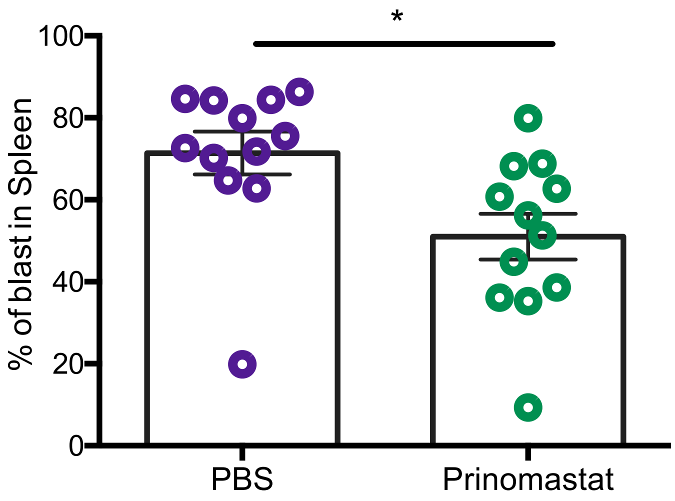


**Figure S2: AML burden in the spleen of PBS- and prinomastat-treated leukemic mice**

AML blast infiltration in the spleen of PBS- and prinomastat-treated leukemic mice. Each dot represents one mouse. *n* = 12 PBS-treated and *n* = 12 prinomastat-treated mice pooled from 2 independent experiments.

Data are shown as mean ± s.e.m. * $p < 0.05$;. *P* values determined using Student's t-test

**Supplementary Video 1: AML induces healthy cell egress from the BM parenchyma**

Representative maximum projection of 3D time-lapse data acquired every 3 minutes for 24 minutes (shown at 3 frames per second) of an area from a leukemic Flk1-GFP mouse (shown in Figure 1A). Healthy haematopoietic cells are shown in red; leukemic blasts are $YFP^+$ and shown in dim green. Ecs are $GFP^+$ and shown in bright green. White arrows indicate the healthy haematopoietic cells intravasating into the circulation.

**Supplementary Video 2: Circulating healthy cells are found as singlets in healthy mice compared to clusters in AML-burdened mice.**

Representative maximum projection of 3D time-lapse data acquired every 3 minutes for 36 minutes (shown at 3 frames per second) of an area from healthy (left) and leukemic (right) Flk1-GFP mouse (shown in Figure 1B). $mTomato^+$ healthy haematopoietic cells are red; leukemic blasts in green. White arrows indicate the clusters of healthy cells in circulation.

**Supplementary Video 3: AML increases vasculature permeability at early disease stages.**

Representative maximum projection of 3D time-lapse data (shown at 3 frames per second) of an area from a healthy (left) and a leukemic (right) Flk1-GFP mouse at early disease stage (blasts in PB <10%), injected with 3mg 65-80kDA TRITC-dextran (shown in Figure 1D). An image was recorded every minute for 10 minutes immediately after injecting the vascular dye in order to assess vascular leakiness. Flk1-GFP$^+$ ECs are shown in green; TRITC-dextran is shown in magenta.

**Supplementary Video 4 and 5: Prinomastat reduces the circulating clusters of healthy cells in AML-burdened mice**

Representative maximum projection of 3D time-lapse data acquired every 3 minutes for 63 minutes (shown at 3 frames per second) of an area from a leukemic PBS-treated mouse (shown in Figure 2B left) and a leukemic prinomastat-treated mouse (shown in Figure 2B right). mTomato$^+$ healthy haematopoietic cells are red; leukemic blasts in green

**Supplementary Video 6: Prinomastat reduces vascular leakiness induced by AML**

Representative maximum projection of 3D time-lapse data (shown at 3 frames per second) of an area from a leukemic PBS-treated (left) and leukemic prinomastat-treated (right) Flk1-GFP mouse at late disease stage (blasts in PB >10%), injected with 3mg 65-80kDA TRITC-dextran (shown in Figure 2D). An image was recorded every minute for 15 minutes immediately after injecting the vascular dye in order to assess vascular leakiness. Flk1-GFP+ ECs are shown in green; TRITC-dextran is shown in magenta.

**Supplementary Video 7: Leukemic blasts demonstrate heterogenous behaviours**

Representative maximum projection of 3D time-lapses acquired every 3 minutes for 102 minutes (shown at 3 frame per second) of an area from a leukemic mouse (shown in Figure 3G). Among leukemic blasts, white arrows highlight the explorative cells. mTomato$^+$ healthy haematopoietic cells are shown in red; leukemic blasts are YFP$^+$ and shown in green.

### Supplementary Materials and Methods

#### Mice

All mice were bred and housed at Imperial College London or Sir Francis Crick Institute.

C57BL/6 wild-type (WT) were purchased from Charles River UK or obtained from the Francis Crick Institute. mT/mG mice[1] were purchased from Jackson Laboratories. Flk1-GFP mice[2] were a gift from Alexander Medvinsky (Unversity of Edinburg). PU1-YFP mice[3] were a gift from Claus Nerlov (University of Oxford). Female mice > 6 weeks old were used.

#### BM chimeras

To generate chimeric mice, whole BM cells were obtained from femurs, tibias and hips of mT/mG donor mice, diluted in PBS and transplanted intravenously into lethally irradiated (two doses of 5.5 Gy separated by 3 hours) WT or Flk1-GFP recipient mice at a dose of 1.5 x $10^6$ cells/mouse. Transplanted mice were kept on baytril-containing water for five weeks following transplantation. >95% chimerism was confirmed after 8 weeks and at that point mice were injected with AML cells and used for intravital imaging experiments.

**AML experimental model**

To follow disease development by both flow cytometry and microscopy, granulocyte-macrophage progenitors (GMPs) were sorted from C57BL/6 wild-type, mT/mG or PU.1-YFP mice. Sorted GMPs were then transfected and transduced with pMSCV-MLL-AF9-GFP-based retroviruses as described by Krivtsov and collegues[4] and transplanted in sub-lethally irradiated mice. >8 weeks post transplantation, recipient mice developed leukaemia characterised by multi-organs infiltration. $GFP^+$ cells were then harvested from BM and spleen and blasts from each primary recipient were labelled as a separate batch and stored. Primary blasts from different mice were thawed, suspended in (PBS) and 100,000 viable cells were transplanted intravenous into secondary, non-conditioned recipient mice. In some experiments, secondary blasts were used. Progressive AML expansion was observed from day 8-10 and full BM infiltration was typically reached between day 20 and 28, with some variability depending on the primary blasts analysed. This was accompanied by infiltration of the spleen and liver, typically delayed compared to BM infiltration.

**Drugs treatment**

For prinomastat (AG3340) treatment, daily 13.5mg/kg prinomastat hydrochloride ≥ 95% (Sigma Aldrich, Cat. # PZ0198) was administered iv from day 7 and continued for 15 days in total. Control mice were injected i.v. with 100 $\mu$l of PBS.

Induction chemotherapy for AML was administered when PB blood infiltration was >15% by injecting 100mg/kg cytarabine (Ara-C) i.v. for 5 days and 3mg/kg doxorubicin (Doxo) for 3 days. Ara-C was co-delivered with Doxo on days 1 to 3 and alone on days

4 and 5 mimic the 7+3 regimen used in patients[5]. Both drugs were obtained from the Imperial College Healthcare NHS trust.

**Flow cytometry**

For hematopoietic and leukemic cell analysis, bones were crushed in PBS with 2 % fetal bovine serum and the cells filtered through a 40 $\mu$m strainer. When spleens were analyzed, the organs were mashed through a 40 $\mu$m strainer using the internal part of 5 or 10ml syringes and resuspended in FACS buffer. For endothelial cells analysis, tibias and femur were crushed, digested with collagenase I (Worthington, UK) at 37° for 20minuteswith 110rpm agitation, and the obtained cells were filtered through a 70 40$\mu$m strainer. The following fluorochrome-conjugated or biotinylated primary antibodies specific to mouse were used: CD3e (145-2C11), CD4 (GK1.5), CD8a(53-6.7), Ter119 (TER119), B220(RA3-6B2), Ly6G(RB-68C5), CD11b(M1/70), c-Kit(2B8), Sca-1(D7), CD150(TC15-12F12.2), CD48(HM48-1), CD45(104) and CD31(MEC13.3) were all from Biolegend. For secondary staining, streptavidin Pacific Orange (Invitrogen) were used. Live and dead cells were distinguished using 4,6-diamidino-2-phenylindole (DAPI, Invitrogen). EdU was detected using the Click-iT EdU kit (Life Technologies). To identify apoptotic cells, an Annexin V kit (BD Pharmingen™ Annexin V APC; BD Biosciences, Cat. # 550474) was used. For the analysis of cellular ROS, CellROX Deep Red Reagent was used, following manufacturer's instructions (ThermoFisher Scientific). Calibrite beads (BD Bioscences) were used to determine the absolute cell number in the population of interest, as described previously[6]. Samples were analysed with a LSR-Fortessa (BD Bioscences) and data were analysed with FlowJo (Tree Star).

**Enzyme-linked immunosorbent assay (ELISA)**

To obtain BM supernatant, tibias and femurs were harvested from healthy control, PBS treated and prinomastat treated AML-burdened mice. With scissors, the metaphysis and diaphysis of long bones were separated and 100$\mu$l of PBS were flushed through diaphysis, collected and re-flushed; cells were then excluded by centrifugation at 400g for 5 min; the supernatant was collected and any remaining cells excluded by centrifugation at 500g for 5 min. BM supernatant was stored at -20° C until used for ELISA. Legend MAX™ Mouse CXCL12 (SDF-1ß) ELISA kit (Biolegend, Cat # 444207), SCF (KITL) Mouse Elisa kit (Thermofisher, Cat. # EMKITL) and Mouse VEGF DuoSet ELISA (R&D system, Cat #DY493) were performed according to the manufacturer's instructions.

**Intravital microscopy**

Intravital microscopy (IVM) was performed using a Zeiss LSM 780 upright confocal microscope equipped with Argon (458, 488 and 514 nm) a diode-pumped solid-state 561 nm laser and a Helium-Neon 633 nm, a tuneable infrared multiphoton laser (Spectraphysics Mai Tai DeepSee 690-1200nm)., 4 non-descanned detectors (NDD) and an internal spectral detector array. Live imaging of the calvarium BM was done as described in previously published reports.[7,8] In some case, healthy hematopoietic cells were labelled in vivo using 10 $\mu$g/ml of CD45.2 antibody in PE or APC (BioLegend). Second harmonic signal was excited at 860-890nm and detected with external detectors. GFP signal: excitation at 488nm with an internal detector, YFP signal: excitation at 488 or 514 nm, internal detectors. mTomato/DsRed and Dextran signals were excited at 561nm and detected using internal detectors. Assessment of vascular leakiness in healthy control, PBS treated and prinomastat treated leukemic mice was carried out by adapting previously published protocols. Briefly, once that the mouse was ready on the microscope stage and positions for analysis selected, 60$\mu$l of low molecular weight TRITC-dextran (3mg/ml, 65-80kDA) was injected (i.v) and continuously recorded (1 frame per minute) for 10 or 15 minutes after injection.

**Image processing and quantification**

Zen black (Zeiss, Germany) software was used to stitch three-dimensional (3D) BM tilescans (tilescans represent individual tiles stitched together to form a composite). ImageJ was used to visualize and process raw data. ImageJ was also used to manually crop out autofluorescent signal from the tissue. For time lapse data, any displacement in the Z-plane caused by artefact movement were corrected by applying four-dimensional (4D) data protocols implemented in ImageJ which allow the registration of the acquired time lapse[9]. Cell and cluster counting were performed manually using the FIJI plugin cell counting while cell tracking was performed semi-automatically using Imaris. Again, time lapses were corrected using the 4D data protocols before importing the selected images to Imaris. Imaris software was used to detect leukaemia cells and create either “surfaces” or “spots” and the semi-automatic cell tracking was done using built-in algorithms that were then manually supervised. Obtained data, including track mean speed, length and linear progression were exported for further analysis.

For leaky vasculature analysis, the TRITC-dextran extravasation in time lapses movies from healthy control, PBS and Prinomastat treated leukemic mice was quantified by measuring the pixel intensity within 3 equally sized and randomly placed region of interest per time frame. The average of these values was subsequently calculated and the fold change increase in intensity for each group plotted in a graph.

**RNA sequencing and analysis**

RNA-sequencing data was downloaded from previously published data (GSE105159)[10] for reanalysis. Read trimming was performed using Cutadapt (v1.9). The trimmed reads were subsequently mapped to the mouse genome (mm10) using HISAT2[11]. *FeatureCounts* from the *Rsubread* package (version 1.34.7)[12,13] with its in-built mm10 annotation was used to obtain gene counts. Genes with a count per million (CPM) in at least 3 samples were included downstream analysis. Count data were normalised using the trimmed mean of M-values (TMM) method and differential gene expression analysis was performed using the limma-voom pipeline (*limma* version 3.40.6)[14]. The R/Bioconductor package *ggplot2* (version 3.2.1) was used to generate the volcano plots. Heatmaps of logCPM were generated using *pheatmap*.

**Human transcriptomic data analysis**

For the analysis, the FPKM values of patients with AML and healthy subjects were taken into consideration. For the latter, four samples of mononuclear cells from bone marrow aspirates obtained from GSE (https://www.ncbi.nlm.nih.gov/geo/query/acc.cgi?acc=GSE61410) were included. The data relating to patients with AML were instead retrieved from the site https://portal.gdc.cancer.gov/ by selecting available cases within TCGA-LAML project. In particular, four representative AML groups established for comparison with the healthy samples, based on genes commonly mutated in adult AML patients[15]:

- *NPM1*-mutated AML group (n=7). Included patients were: TCGA-AB-2861, TCGA-AB-2869, TCGA-AB-2871, TCGA-AB-2877, TCGA-AB-2925, TCGA-AB-2931, TCGA-AB-2932.
- *FLT3*-mutated AML group (n=5). Included patients were the following: TCGA-AB-2811, TCGA-AB-2814, TCGA-AB-2851, TCGA-AB-2873, TCGA-AB-2910.
- *NPM1*/*FLT3*-mutated AML group (n=5). Included patients were: TCGA-AB-2818, TCGA-AB-2895, TCGA-AB-2900, TCGA-AB-2919, TCGA-AB-2924.
- "Other AML subtypes" group (n=7). This group included patients carrying mutations within genes, others than *FLT3*, *NPM1* and *MLL*, recurrently altered in adult AML. The latter were *DNMT3A*, *RUNX1*, *CEBPA*, *IDH2*. Included patients were: TCGA-AB-2845, TCGA-AB-2884, TCGA-AB-2890, TCGA-AB-2927, TCGA-AB-2936, TCGA-AB-2949, TCGA-AB-2959.

An additional group (*MLLT3-KMT2A* rearranged AML) was instead built up on RPKM data from Tyner et al[16]. Patients of interest (n=8) were: 13-00353, 13-00513, 14-00015, 14-00434, 14-00464, 14-00504, 15-00296, 15-00633.

.

Fold Change was calculated as the base two logarithm of the ratio between the means of the two groups, placing the control, or the healthy ones, in the denominator

#### Quantification and statistical analysis

Raw data was visualized and processed using Microsoft Excel and GraphPad Prism (GraphPad Software Inc.). Group means were compared using the unpaired Student's t test. For multiple comparison, one-way ANOVA with post hoc Tukey test or Bonferroni correction or multiple t-test with Holm-Sidak correction was used. For all data, differences were considered significant for p<0.05*, **p<0.01 :***p<0.001; ****p<0.0001. Number of used animals, together with other statistical details, can be found in the figure legends.

#### References

1. Muzumdar MD, Tasic B, Miyamichi K, Li L, Luo L. A global double-fluorescent Cre reporter mouse. *Genesis*. 2007;45(9):593-605.

2. Xu Y, et al. Neuropilin-2 mediates VEGF-C–induced lymphatic sprouting together with VEGFR3. *J Cell Biology*. 2010;188(1):115-130.

3. Kirstetter P, Anderson K, Porse BT, Jacobsen SEW, Nerlov C. Activation of the canonical Wnt pathway leads to loss of hematopoietic stem cell repopulation and multilineage differentiation block. *Nat Immunol*. 2006;7(10):1048-1056.

4. Krivtsov AV, et al. Transformation from committed progenitor to leukaemia stem cell initiated by MLL–AF9. *Nature*. 2006;442(7104):818-822.

5. Wunderlich M, et al. AML cells are differentially sensitive to chemotherapy treatment in a human xenograft model. *Blood*. 2013;121(12):e90-e97.

6. Hawkins ED, Hommel M, Turner ML, Battye FL, Markham JF, Hodgkin PD. Measuring lymphocyte proliferation, survival and differentiation using CFSE time-series data. *Nat Protoc*. 2007;2(9):2057-2067.

7. Hawkins ED, et al. T-cell acute leukaemia exhibits dynamic interactions with bone marrow microenvironments. *Nature*. 2016;538(7626):518-522.

8. Rashidi NM, et al. In vivo time-lapse imaging shows diverse niche engagement by quiescent and naturally activated hematopoietic stem cells. *Blood*. 2014;124(1):79-83.

9. Preibisch S, Saalfeld S, Schindelin J, Tomancak P. Software for bead-based registration of selective plane illumination microscopy data. *Nat Methods*. 2010;7(6):418-419.

10. Duarte D, et al. Inhibition of Endosteal Vascular Niche Remodeling Rescues Hematopoietic Stem Cell Loss in AML. *Cell Stem Cell*. 2017;22(1):64-77.e6.

11. Kim D, Paggi JM, Park C, Bennett C, Salzberg SL. Graph-based genome alignment and genotyping with HISAT2 and HISAT-genotype. *Nat Biotechnol*. 2019;37(8):907-915.

12. Liao Y, Smyth GK, Shi W. featureCounts: an efficient general purpose program for assigning sequence reads to genomic features. *Bioinform Oxf Engl*. 2013;30(7):923-930.

13. Liao Y, Smyth GK, Shi W. The R package Rsubread is easier, faster, cheaper and better for alignment and quantification of RNA sequencing reads. *Nucleic Acids Res*. 2019;47(8):gkz114-.

14. Lord BI, Testa NG, Hendry JH. The relative spatial distributions of CFUs and CFUc in the normal mouse femur. *Blood*. 1975;46(1):65-72.

15. Döhner H, et al. Diagnosis and management of AML in adults: 2017 ELN recommendations from an international expert panel. *Blood*. 2017;129(4):424-447.

16. Tyner JW, et al. Functional genomic landscape of acute myeloid leukaemia. *Nature*. 2018;562(7728):526-531.
